# Earthquake-induced habitat migration in a riparian spawning fish: implications for conservation management

**DOI:** 10.1101/229872

**Authors:** Shane Orchard, Michael J. H. Hickford, David R. Schiel

## Abstract

*Galaxias maculatus* is a riparian spawning fish that supports an important recreational fishery in New Zealand with spawning habitat requirements strongly structured by salinity gradients at rivermouths. This study reports changes to the spawning habitat following a series of large earthquakes that resulted in widespread deformation of ground surfaces in the vicinity of waterways. Assessments of habitat recovery focussed on two rivers systems, the Avon and Heathcote, with pre-disturbance data available over a 20 year period. Recovery dynamics were assessed by field survey and mapping of spawning habitat prior to and on seven occasions after the disturbance event. Riparian land-use and management patterns were mapped and analysed using overlay methods in GIS. Habitat migration of up to 2 km occurred in comparison to all previous records and several anthropogenic land uses have become threats due to changed patterns of co-occurrence. Incompatible activities now affect more than half of the spawning habitat in both rivers, particularly in areas managed for flood control purposes and recreational use. The results are an example of landscape scale responses to salinity and water level changes driven by tectonic dynamics. These dynamics are not the source of the stress *per se*, rather, they have increased exposure to pre-existing stressors. The case illustrates important principles for managing subtle, yet widespread, change. Adaptive conservation methods and investments in information are priorities for avoiding management failure following environmental change.

## Introduction

### Earthquake recovery context

The Canterbury region of New Zealand was affected by a sequence of major earthquakes in 2010 and 2011. The most devastating of these was a M_w_ 6.3 earthquake centred beneath the city of Christchurch that caused widespread damage and loss of life (Quigley et al., 2016). After six years of recovery activities the process has entered a more strategic phase. The focus is now on longer term adaptation to environmental and societal change. Important land-use decisions remain for many geographical areas and with regards to many aspects of the natural and built environment. Examples relevant to waterway management include responses to water quality, erosion, flood risk and coastal inundation issues, and the potential re-zoning of large tracts of floodplain and riparian land. Existing statutory arrangements apply to many of the recovery planning requirements and identify institutional responsibilities. Due to the scale and impact of the event bespoke legislation was created to facilitate recovery. Organisations involved include new planning entities with specific tasks (Regenerate Christchurch, 2017) and a wide range of interests across central, regional, and local government, non-governmental organisations, and local community groups.

Initially, urgent decisions were made to address risks to property and life, and to reinstate essential infrastructure. Remaining decisions have the benefit of more time. There is a unique opportunity to secure benefits through earthquake recovery planning in relation to historical degradation of natural environments and improved resilience to future events. Natural values in the affected areas have thus far received less attention, but include traditional cultural uses such as the wild harvest of food and fibre (Jolly & Ngā Papatipu Rünanga Working Group, 2013; Lang et al., 2012), risk reduction functions (Orchard, 2014), and habitat for many indigenous and migratory species with protected status. However, knowledge gaps are a barrier to securing benefits through the planning process. Information requirements include quantifying impacts of the earthquakes and identifying opportunities for future gains.

### Spawning habitat of īnanga

In the present study, our particular focus is *Galaxias maculatus*, or ‘īnanga’, a riparian spawning fish. *G. maculatus* is an amphidromous species currently listed as ‘at risk - declining’ in the New Zealand Threat Classification System (Goodman et al., 2014). Reversing the decline of īnanga is addressed in many statutory documents as well as non-statutory plans and is a priority issue for Māori. Juvenile fish are harvested in an iconic recreational and culturally important fishery (McDowall, 1984). The harvest of īnanga and other ‘whitebait’ species creates an ongoing tension between conservation and sustainable use. However, use and non-use interests share the objective of enhancing īnanga populations. The protection of spawning habitat is an urgent and practical goal due to a history of degradation associated with land-use changes near lowland waterways (McDowall, 1992; McDowall & Charteris, 2006).

Īnanga has a specialised reproductive strategy that is synchronised with the spring tide cycle which strongly influences the distribution of spawning sites (Burnet, 1965). Spawning sites occur close to the maximum upstream extent of saltwater intrusion and occupy only a narrow elevation range (Taylor, 2002). Eggs are laid in riparian vegetation just below the spring tide high-water mark and hatch in response to inundation after a 2-4 week development period (Benzie, 1968b). The composition and condition of riparian margins at these specific sites is critical to spawning success (Hickford & Schiel, 2011a).

This specificity suggested that earthquake-induced land deformation could affect habitat in several ways. First, disturbance could reduce the availability or condition of existing spawning sites, and enduring changes might result from vegetation recovery effects. Second, large scale impacts were possible due to physicochemical effects. This was the particular focus of our study in light of suspected earthquake-driven hydrodynamic changes and the reported structuring of habitat by salinity (Richardson & Taylor, 2002; Taylor, 2002). Because there was no prior salinity baseline available, our focus was direct detection of changes in the distribution of spawning sites. By reconstructing a spawning site distribution baseline using data from previous studies, this comparison was possible for the consideration of earthquake effects. The objectives of the study were therefore to quantify the pre-and post-quake spawning site distribution against riparian land uses and evaluate distributional effects to identify management implications.

## Methods

### Study area

The two study catchments are the Avon River (Ōtākaro) and Heathcote River (Ōpāwaho) (Figure 1). These are spring-fed, lowland waterways with small average base flows (approx. 2 and 1 cumecs respectively) originating within the city of Christchurch, New Zealand (White et al., 2007). The catchments are heavily urbanised, particularly in their upstream reaches. The two waterways are often channelized through the use of bank stabilisation engineering and flow regulation structures including flood-gates. The lower catchments are characterised by extensive wetlands and saltmarsh areas that comprise the Avon-Heathcote Estuary / Ihutai (Figure 1). These are remnants of a larger and relatively mobile ecosystem of coastal hydrological features (Kirk, 1979).

**Figure 1.**
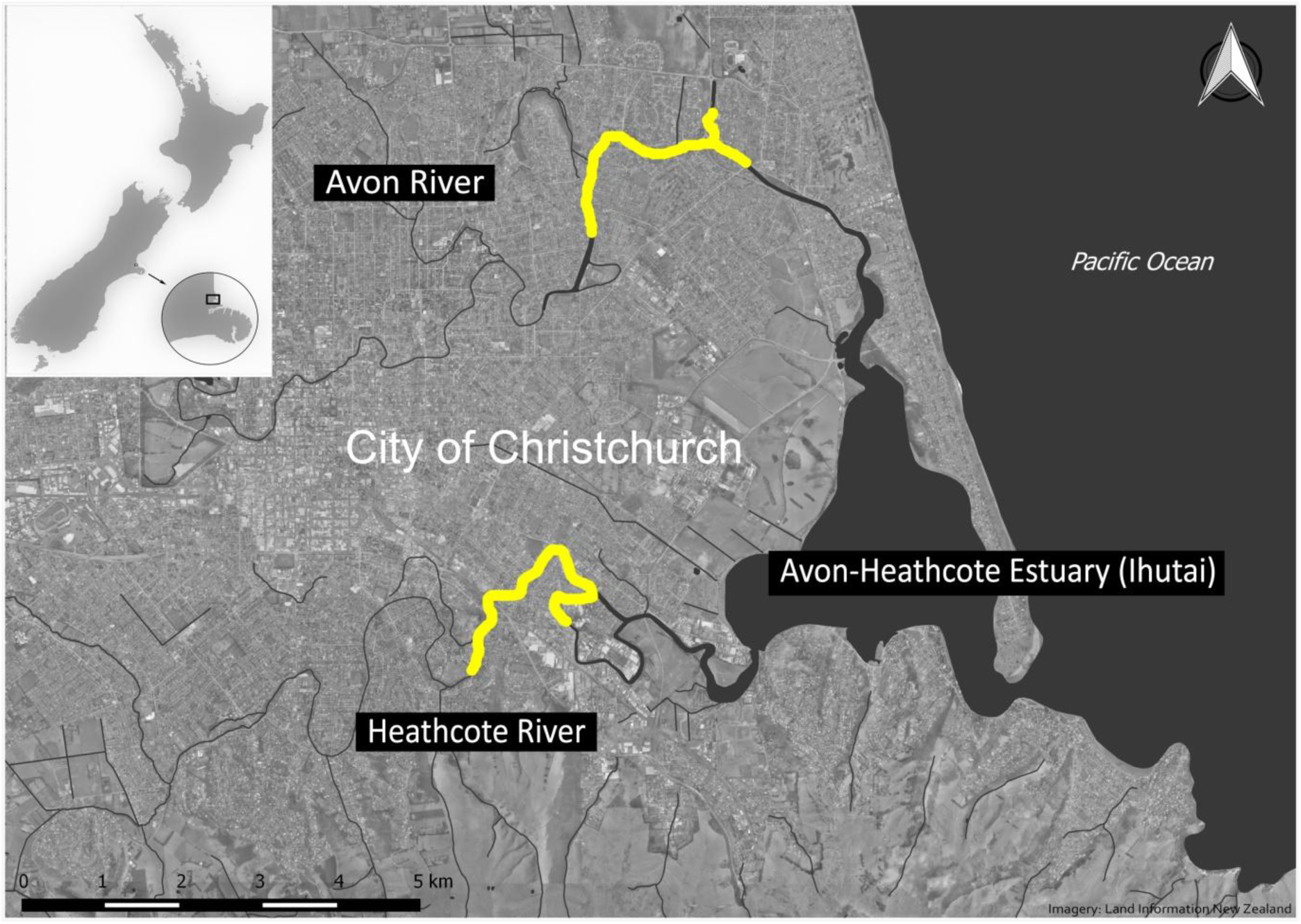
Location of the Avon and Heathcote Rivers in Christchurch, New Zealand, showing reaches surveyed for īnanga spawning in the post-quake studies (in yellow).

Vertical seismic shifts and lateral spread were pronounced in the vicinity of Christchurch waterways particularly towards the estuary (Hughes et al., 2015). Changes in ground levels in and around the estuary were in the order of ± 0.5 m with a trend towards uplift in the south and subsidence in the north (Beaven et al., 2012). Hydrodynamic modelling of the estuary showed extensive bathymetric change and an estimated 15% reduction in the estuarine tidal prism (Measures et al., 2011).

### Pre-earthquake baseline

A literature review was completed to identify pre-quake spawning records augmented with information from current researchers (M. Taylor, S. McMurtrie, C. Meurk, pers. comm.) and records from the National Īnanga Spawning Database (www.inangaconservation.nz). Historical spawning site data were restricted to sites identified through the observation of eggs in riparian vegetation. All information was digitised in GIS by identifying coordinates for upstream and downstream extents from maps, photographs or co-ordinates provided in the original reports. Where this information was not available, locations were estimated using the text descriptions provided. Semi-continuous stretches of spawning were lumped into a single reach in some cases, generally following the description given in the original records.

### Post-earthquake studies

A census-style survey methodology was used with the objective of detecting all spawning occurrences at the catchment scale following the methods of Orchard & Hickford (2017). The search areas were approximately 4 km reaches in each river (Figure 1). The survey area extended from the downstream transition to saltmarsh vegetation, which is unsuitable for spawning (Mitchell & Eldon, 1991), to 500 m upstream of the inland limit of saltwater. In the Avon this included the confluence with a prominent tributary to the north. The saltwater limit was established using conductivity/temperature loggers (Odyssey, Dataflow Systems Ltd, NZ) deployed during spring tide sequences and additional spot measurements using a handheld conductivity/salinity/temperature meter (YSI Model 30, YSI Inc., USA). The survey period included the peak spawning months (Taylor, 2002) over two years. Surveys commenced five days after the peak tide in the spring tide sequence and followed a set schedule to minimise temporal confounding effects between months (Table S1). Reaches surveyed later in the schedule were more sensitive to egg mortality effects due to the time elapsed since spawning. Results are more likely to underestimate the extent of spawning occurrences in these areas, but are comparable between months.

The search area was surveyed systematically in the first two months of the study by conducting three searches for eggs within contiguous 5 m blocks along each riverbank. Each search involved opening up the vegetation down to ground level at random locations within the block following a transect line perpendicular to and spanning the high water mark. On subsequent months, the survey effort was reduced to areas of potential habitat following a habitat classification system (Orchard & Hickford, 2017). Whenever eggs were found, the survey was extended 50 m either side of the last occurrence to confirm the full extent of the spawning site. Spawning sites were defined as the area occupied by continuous or semi-continuous patches of eggs. Upstream and downstream extents were established and the width of the egg band measured on the centreline of the search transects within the extent of the site (minimum three). Zero counts were recorded where these occurred such as when the egg patch was not continuous. Area of occupancy (AOO) was calculated as length × mean width. The total number of eggs present was calculated by sub-sampling patches. At each width measurement location, eggs were counted in a 10 × 10 cm quadrat placed in the centre of the egg band. Productivity was calculated as mean egg density × AOO.

Riparian land uses and management activities were mapped in the field using 0.075 m resolution post-quake aerial photographs (Land Information New Zealand, 2016). Anthropogenic stressors were identified based on reported incompatibility with īnanga spawning sites (Hickford & Schiel, 2011a, 2011b; Mitchell, 1994). Areas affected were delineated using aerial photographs in the field and digitised for overlay analysis in QGIS v2.8.18 (QGIS Development Team, 2016). Four classes of land use activities were classified as threats to spawning habitat. These were bank stabilisation using engineered structures, invasive species control, mowing of recreation reserves, and vegetation removal for flood management. Threats from riverbank engineering were defined on the basis of surfaces devoid of any vegetation capable of supporting spawning (Mitchell, 1994). Examples include retaining walls, bridge abutments, riprap, and other bank stabilisation works. Invasive species control was classed as a threat where it involved spraying or extensive mechanical clearance (e.g., using scrub cutters, line trimmers & similar). This recognises that vegetation suitable for spawning may take several months to recovery following clearance activities (Hickford & Schiel, 2014). Mowing was classed as a threat where it resulted in short grass conditions at the top of the riverbank in the location of spawning habitat.

## Results

### Pre-earthquake spawning distribution

Eighteen pre-quake spawning studies spanning a 25 year period were identified, most of which involved surveys in both catchments. Thirteen of these had quantified spawning in the Avon and nine in the Heathcote (Table 1). In some years field surveys were conducted that did not find any spawning and these records are not shown in Table 1. In the Avon, most of the spawning occurrences have been in the Avondale Road area and often found a short distance upstream from the road bridge on the true right (Table 1). The maximum extent of pre-quake spawning sites recorded in any one year was 2000 m. This also represents the maximum extent of the spawning reach based on all known records.

**Table 1.**
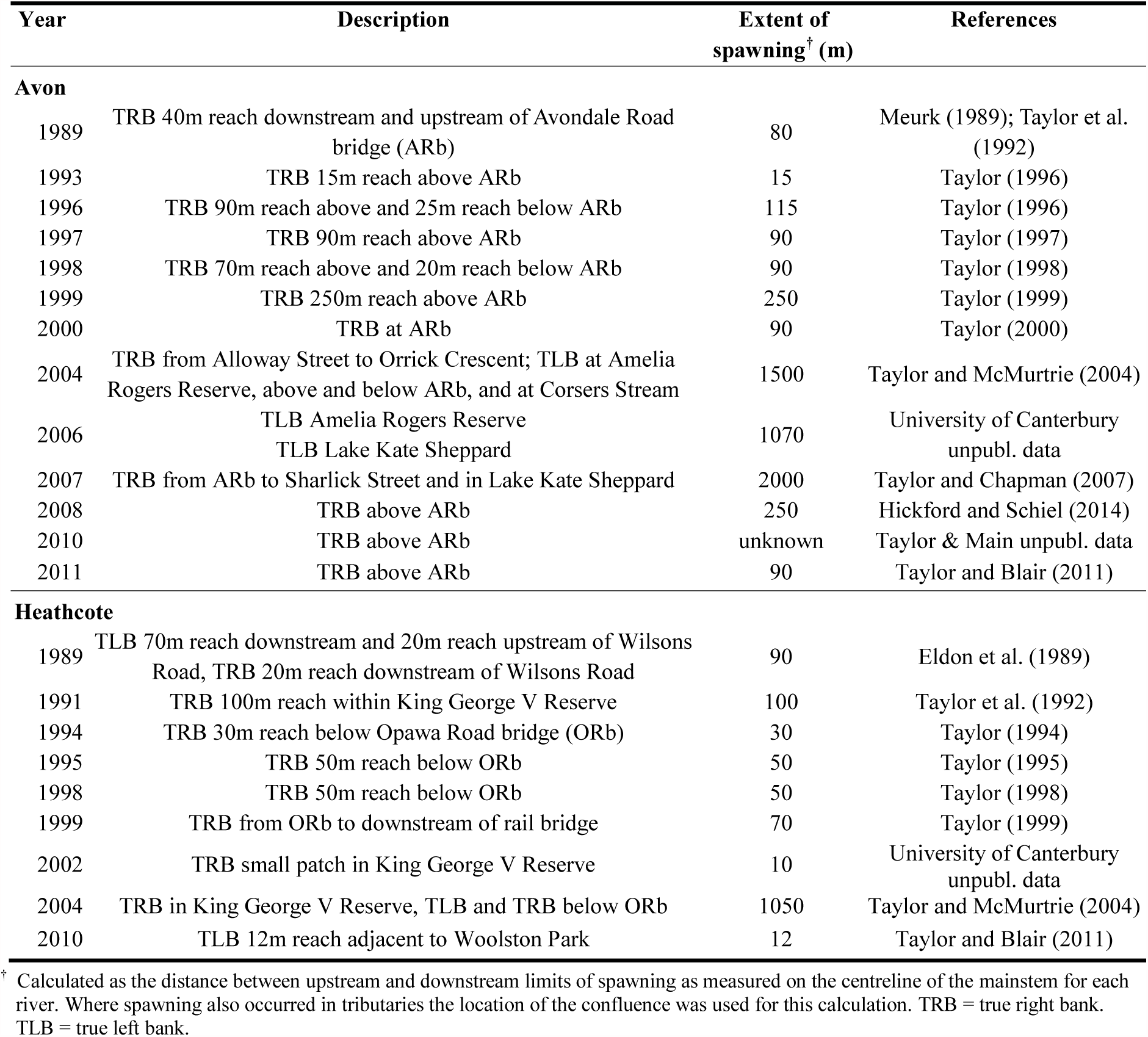
Extent of Īnanga spawning habitat utilised in the Avon and Heathcote Rivers over the period 1989–2014 from all known records.

In the Heathcote, most of the records have been in the vicinity of Opawa Road (Table 1). Although the downstream limit of all records is c. 1 km further downstream this relates to only two observations of spawning below Opawa Road in the 25 year period. However, the first spawning recorded in the catchment was much further upstream (> 3 km). At the time the river was under the influence of a floodway, constructed in 1986, that effectively shortened the length of the river. In 1994 a tidal barrage was installed to reduced saline intrusion and this resulted in a shift of c. 2km downstream in the upstream limit of spawning (Taylor & McMurtrie, 2004). These variations in the location of pre-quake sites complicate analyses of the extent of spawning habitat based on pooled records. However, the maximum extent of pre-quake spawning recorded in any one year was 1050 m in 2004 (Table 1).

### Post-quake studies

#### Spawning extent

A total of 85 spawning sites were identified in the 2015 post-quake survey. These were distributed along 2.4 km of riverbank in the Avon and 2.5 km in the Heathcote. In both rivers there were marked differences in the spawning distribution in comparison to previous records (Figure 2). In the Avon, the spawning reach had expanded approximately 250 m upstream and 180 m downstream of the previous extent. In the Heathcote, the changes were more pronounced with spawning recorded 1.5 km downstream of all previous records ((Figure 2a)). The 2016 survey identified a total of 101 spawning sites, some of which represented repeat use of 2015 sites. In the Avon, the upstream and downstream limits were very close to those recorded in 2015. In the Heathcote, the upstream limit was also similar to 2015, but the spawning reach extended a further 400 m downstream ((Figure 2b)).

**Figure 2.**
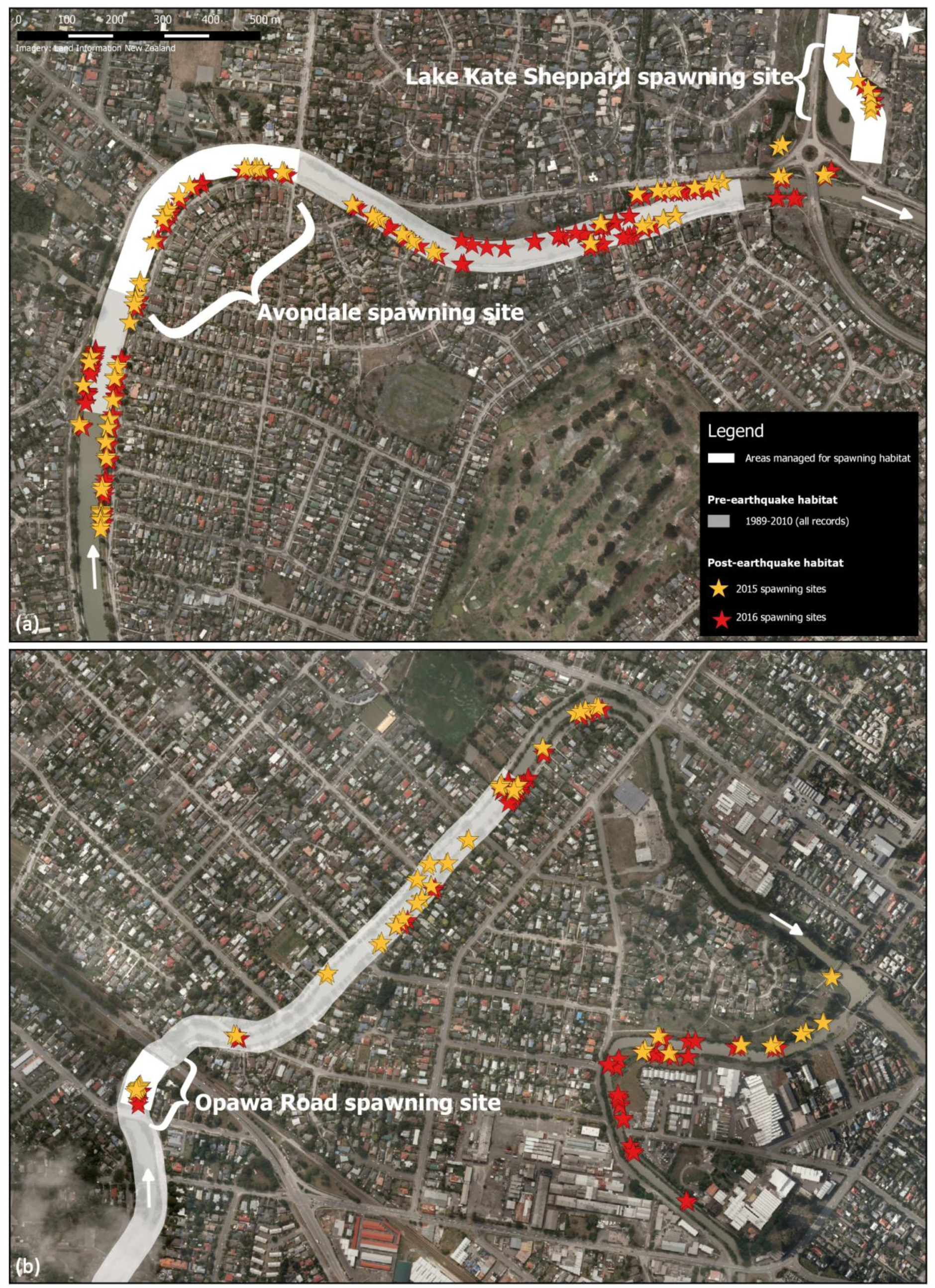
Post-quake īnanga spawning distribution overlaid on the maximum pre-quake extent of spawning from all known records. Well-known pre-quake spawning sites are shown. (a) Avon River (Ōtākaro). (b) Heathcote River (Ōpāwaho).

#### Distribution of threats and protected areas

There are three areas managed to protect spawning habitat at well-known sites (Figure 3). The protection mechanisms include recognition in local authority plans and implementation of compatible riparian management on the ground. There is also a considerable reach in the lower Heathcote that is not subject to vegetation clearance for flood or reserves management purposes primarily due to being a neglected part of the river for maintenance. Part of this reach is characterised by tall woody riverbank vegetation and the remainder is downstream of the tidal barrage where there is less need for flood management-related channel maintenance ((Figure 3b)).

**Figure 3.**
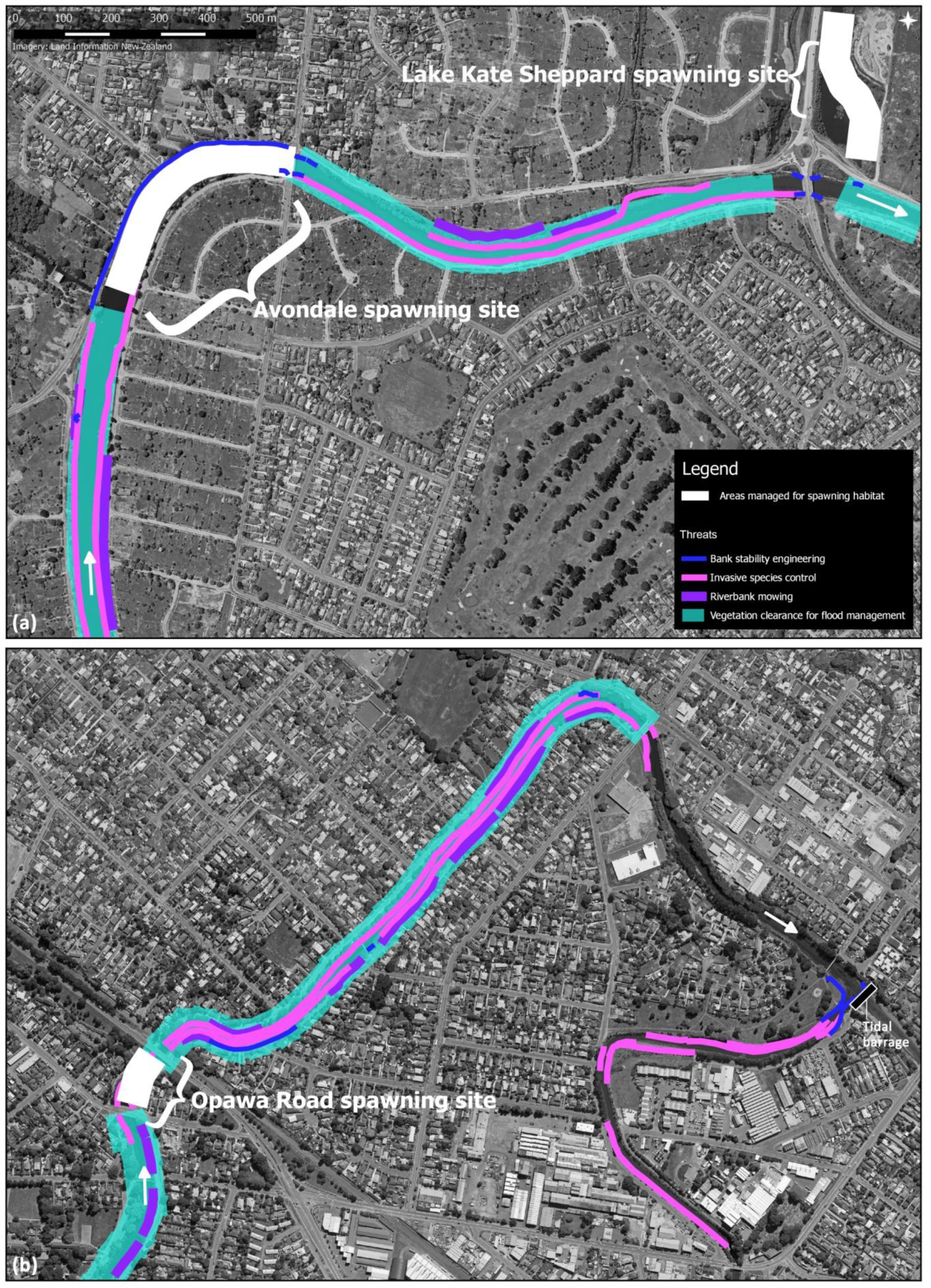
Distribution of post-quake anthropogenic threats associated with riparian land uses and management activities in the study area. (a) Avon River. (b) Heathcote River.

Threats from riverbank engineering occupied only a small proportion of the post-quake spawning extent in each river (Figure 3). Extensive channelization using gravel embankments is also found in the Avon. Although the area available for spawning may be reduced by these structures, they were not classified as threats based on observations of spawning if suitable vegetation co-occurred. Invasive plant species that have historically been the subject of spraying or mechanical clearance are widespread in the study area. In the Avon the major concern is Yellow Flag Iris (*Iris pseudacorus*). It is distributed throughout the spawning reach with the exception of sections engineered with gabion baskets and in Lake Kate Sheppard. This species is largely absent from the Heathcote and instead Reed Canary Grass (*Phalaris arundinacea*) is the major concern and is the dominant canopy species in many areas. In addition, *Glyceria maxima* and *Rubus fruticosus* are present there. There were no major eradication campaigns during the study period despite the severe level of infestation. With the exception of the protected areas (as above), vegetation control for flood management was conducted throughout the study area on a semi-regular basis consistent with previous years. This involves clearance of all bank vegetation using scrub cutters or line trimmers. Riparian mowing occurs in discrete areas in both river systems associated with a variety of parks and reserves in the river corridor (Figure 3).

#### Area of occupancy of egg production

In 2015, the total area of occupancy (AOO) of spawning habitat was 152.5 m^2^ in the Avon and 75.4 m^2^ in the Heathcote based on maximum figures recorded at each site across all four surveys. Total egg production was 11.8 million eggs (Avon 6.9 × 10^6^, Heathcote 4.9 × 10^6^). In 2016, egg production was higher (Avon 13.9 × 10^6^, Heathcote 5.0 × 10^6^) despite the survey period being reduced to only three months. The AOO was also higher in both rivers (Avon 472.9 m^2^, Heathcote 99.1 m^2^) although average egg densities were lower. The marked increase in AOO in the Avon was associated with several new large spawning sites that were not utilised in 2015 in addition to re-use of other sites. In both years, AOO and productivity were not evenly distributed across the study area. High production was not always correlated with AOO due to differences in egg densities (Figure 4). Egg densities of >10 eggs cm^−2^ were recorded at several sites with the highest being 13.5 cm^−2^.

**Figure 4.**
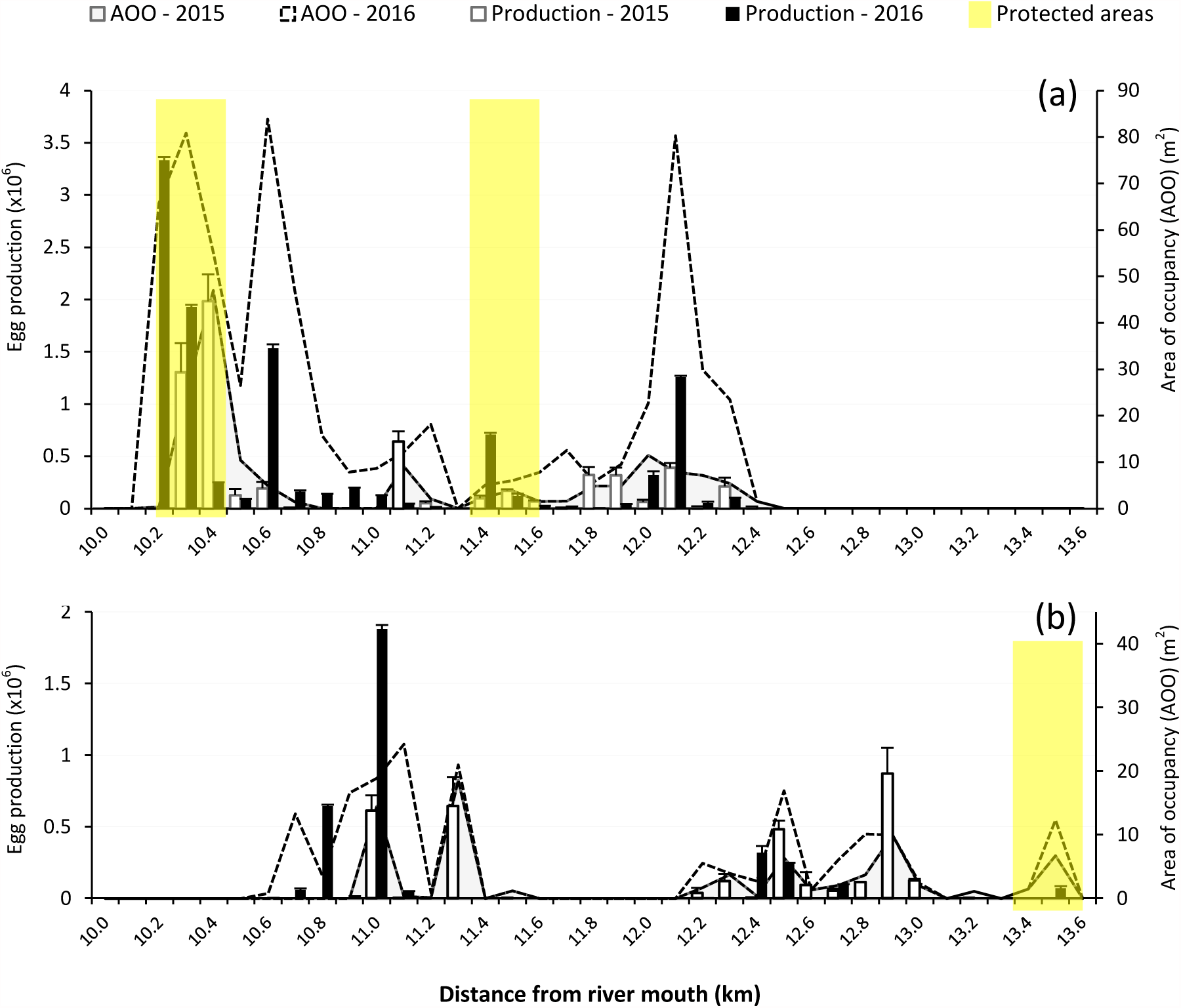
Post-quake area of occupancy (AOO) and productivity of īnanga spawning sites presented as aggregated data for contiguous 100 m reaches of the Avon and Heathcote Rivers. Egg production is shown as the total recorded in the surveys conducted each year (n=4, 2015; n=3, 2016). AOO is presented as the maximum area occupied by spawning sites each year. River kilometres are measured from the entrance into the Avon-Heathcote Estuary / Ihutai following the main channel lines for each river. Error bars are standard errors of the mean.

### Effectiveness of protected areas

In the Avon, the proportion of the AOO occurring in protected areas was 70% in 2015. In 2016 this figure had decreased to only 28% reflecting many new sites discovered in other locations. In the Heathcote, the proportion of AOO protected was very low (11% and 6% for the two years respectively) reflecting that the majority of spawning sites were discovered at sites never previously known for spawning. Post-quake egg production was considerable in the unprotected areas (Figure 5). In the Avon, the proportion of egg production outside the protected areas was 28% in 2015 and 38% in 2016 ((Figure 5a)).In the Heathcote, 82% of egg production occurred outside of the protected areas in 2015 and 98% in 2016 ((Figure 5b)). On average across the two years of post-quake studies, only 4.5% of the spawning reach was protected in the Heathcote and 27.6% in the Avon (Figure 6). Vegetation clearance for reserves and flood management purposes was observed at many of the unprotected spawning sites after egg deposition had occurred (see Supplementary Material). Repeat egg surveys at some of these sites after the vegetation clearance indicated close to 100% egg mortality. This is consistent with previous studies (Hickford & Schiel, 2014).

**Figure 5.**
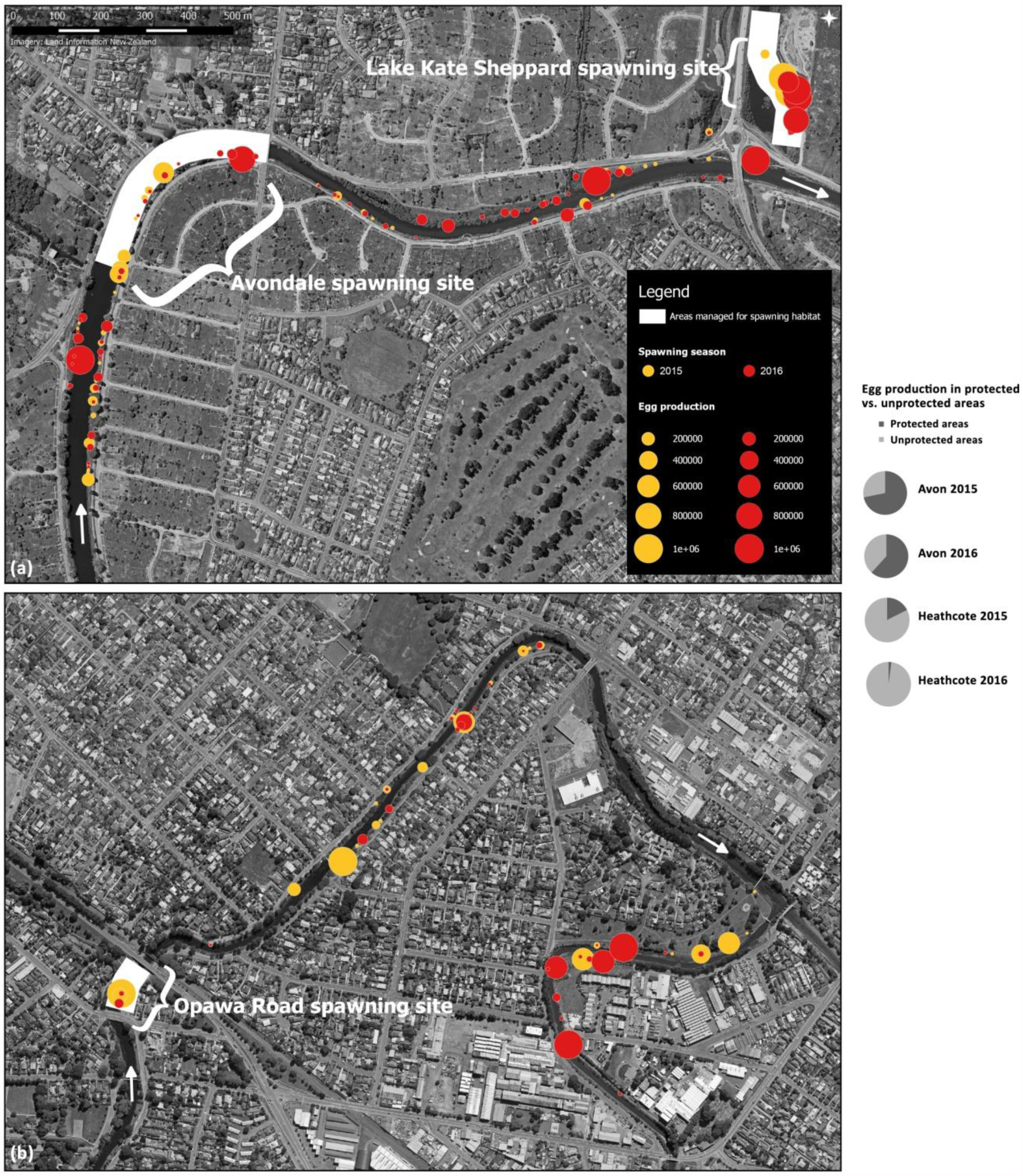
Spatial distribution of post-quake īnanga egg production and proportion of post-quake production that occurred in protected areas. (a) Avon River (Ōtākaro). (b) Heathcote River (Ōpāwaho).

**Figure 6.**
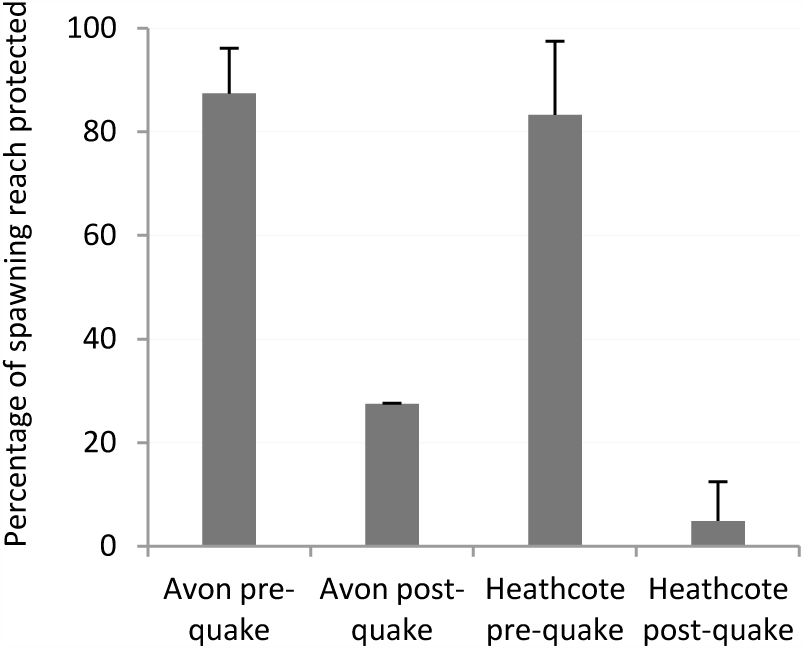
Percentage of the known Īnanga spawning reach in protected areas in the Avon and Heathcote rivers before and after the 2010–11 Canterbury earthquake sequence. Error bars are standard errors of the mean for the periods.

## Discussion

### Evidence for habitat migration

The Christchurch waterways have one of the most extensive records of Īnanga spawning site surveys for any catchment in New Zealand (Taylor, 2002). However, variability in the search effort and methodologies used in the historical surveys are among the sources of uncertainty in characterising the pre-quake baseline. Data used were restricted to confirmed spawning site locations as indicated by the observation of eggs in riparian vegetation. Records of shoaling adult fish in the spawning season are also present in the national database and technical reports and often associated with the location of spawning. These records were considered to be unreliable for quantifying spawning habitat due the mobility of shoals and unknown timing of spawning. In the Avon, the majority of historical spawning has been recorded at the Avondale site. In this vicinity the extent of spawning reach steadily increased since discovery of the site in 1989 with assistance from protection from mowing (Taylor, 1999). In 2004, new sites were identified further downstream in the mainstem, and in 2006 spawning was found at Lake Kate Sheppard and then regularly thereafter ((Figure 2a)). This is an area of restored riparian margins in a tributary waterway and lake system located close to the mainstem. In the Heathcote, the pre-quake distribution has shifted by up the 3 km over the 25 years for which records are available. This has been associated with engineering of the floodway (Taylor, 1998). However, spawning has centred on the Opawa Road site since 2004. Only two sites have been recorded further downstream in all known records. Earthquake-induced migration of habitat a further 1.5 km downstream in 2015 and 1.9 km in 2016 represents a major change in spawning habitat distribution.

### Effectiveness of protected areas

A high proportion of Īnanga spawning now occurs outside of the areas established for protection. Risk exposure is now greater due to the co-concurrence of habitat with anthropogenic threats. Earthquake-induced change is not the source of heightened vulnerability *per se*. Rather, this is an effect of natural dynamics that have increased exposure to pre-existing stressors. These activities are now threats that require management to achieve conservation objectives. Mowing of vegetation within riparian reserves co-occurs with several spawning sites in both river systems. It is a particular issue where the spring high tide water levels are sufficient to inundate riparian terraces. These provide locations where spawning habitat may be relatively expansive in comparison to areas with steeper topography. Vegetation clearance using scrub bars also occurs on the bank face throughout the study area for flood management purposes with the exception of locations specifically managed for Īnanga spawning, being the three established pre-quake spawning sites (Figure 4). Compared to reserve maintenance activities, vegetation clearance for flood management affects the upper intertidal zone of the waterway margin. At many locations this results in a direct overlap with the spawning habitat elevation band. High egg mortality from mowing and grazing has been previously reported (Hickford & Schiel, 2014). This is believed to be mostly attributable to UV irradiation or the drying out of eggs (Hickford et al., 2010; Hickford & Schiel, 2011b). Recovery from vegetation clearance can take many months, with the re-establishment of sufficient cover being a critical factor (Hickford & Schiel, 2014). In addition, these activities may occur after eggs have been laid in vegetation that would otherwise have been suitable for spawning. This was observed at many of the spawning sites recorded in this study and is particularly problematic for conservation. Due to the gregarious behavioural ecology of *G. maculatus* (Benzie, 1968a; McDowall, 1990), the majority of spawning production is typically supported by only a few sites in the catchment in each spawning event. This contributes to the vulnerability of spawning to stochastic events. Anthropogenic threats affecting these highly productive sites may have a large impact on the total egg production on a seasonal basis.

### Lessons for adaptive management

This case illustrates important principles for managing subtle yet widespread change. Habitat migration was not detected by conservation management practitioners. Pre-disturbance land-use activities had continued without adaptation exposing the habitat to increased risk despite its apparent expansion. Adaptive management responses are needed to control anthropogenic stressors in areas that have now become Īnanga spawning habitat. Achieving this requires further work to develop solutions that accommodate other necessary or desirable activities in the riparian zone. Although historical AOO figures are not available, the post-quake studies indicate that in both catchments the extent of spawning habitat is now greater than all previous records. This is a positive finding and suggests a potential improvement in the opportunities available for accommodating incompatible activities through tools such as spatial planning. If these are addressed and solutions identified, conservation gains could be secured in terms of increasing the area of protecting habitat and ultimately improved egg production.

Implementation of statutory protection adds another dimension to this case. It is important to note that protection of the post-quake habitat is a legislative requirement. However, conservation policy frequently suffers from implementation gaps in practice (Knight et al., 2008), often resulting from a lack of attention to methods that are effective in the societal context (Knight et al., 2010). Dynamic environments and spatio-temporal variation create additional challenges for the design of effective methods. Our results illustrate that investments in information are a pivotal activity for achieving this in practice. Regular monitoring or predictive modelling could provide solutions for evaluating change, but they must be coupled with appropriate responses to facilitate adaptive approaches.

Lastly, the effects described here are an example of landscape-scale responses to infrequent tectonic dynamics. They have likely been mediated by hydrological and salinity changes together with smaller-scale effects on ground surfaces in the riparian zone. In the Heathcote, the magnitude of horizontal shift suggests that salinity effects may be involved and this deserves further investigation. Despite these unknowns, the opportunity for learning is clear. Post-earthquake studies present opportunities to evaluate many aspects of socio-ecological systems for impacts and associated responses. Not only are tectonic events relatively common in evolutionary time, they may exert similar effects to climate change through influencing water levels and salinity gradients relative to existing topography (Beavan & Litchfield, 2012). Earthquakes present unique and important opportunities to study vulnerable ecosystems and provide examples of real-life adaptation in action. In turn, this may assist in developing methods to achieve conservation objectives and avoid implementation failures in the face of ongoing change.

## Acknowledgements

We thank Mark Taylor, Shelley McMurtrie and Colin Meurk for providing historical records. We acknowledge the many volunteers and staff of the Waterways Centre for Freshwater Research and Marine Ecology Research Group who assisted with the post-quake field studies, and local government staff for information on riparian management activities. Funding was provided by the Ngāi Tahu Research Centre, Institute of Professional Engineers of New Zealand Rivers Group, Brian Mason Scientific and Technical Trust, and a New Zealand Ministry of Business, Innovation and Employment grant (C01X1002) in conjunction with the National Institute of Water and Atmospheric Research.

## Supplementary Material

**Table S1.** Survey periods and tidal cycle data for post-earthquake *G. maculatus* spawning surveys

| Year | Month of spawning | Peak tidal cycle start | Peak tidal cycle end | Peak tidal height* (m) | Survey period |  |
| --- | --- | --- | --- | --- | --- | --- |
|  |  |  |  |  | Heathcote | Avon |
| 2015 | February | Feb 22 | Feb 25 | 2.6 | Mar 3-6 | Mar 7 -15 |
| 2015 | March | Mar 20 | Mar 23 | 2.6 | Mar 29 -<br>Apr 3 | Apr 4-11 |
| 2015 | April | Apr 18 | Apr 20 | 2.6 | Apr 26-30 | May 1-8 |
| 2015 | May | May 17 | May 19 | 2.6 | May 26-30 | Jun 1-6 |
| 2016 | February | Feb 10 | Feb 14 | 2.5 | Feb18-22 | Feb 23-29 |
| 2016 | March | Mar 10 | Mar 13 | 2.6 | Mar18-22 | Mar23-97 |
| 2016 | April | Apr 7 | Apr 11 | 2.6 | Apr14-18 | Apr19-26 |

\* predicted tide levels above Chart Datum at Port of Lyttelton (Lat. 43° 36' S Long. 172° 43' E) (Source: Land Information New Zealand).

**Table S2.** Habitat quality classes.

| Class | Quality of habitat for supporting spawning | Expected egg mortality rate | Criteria |
| --- | --- | --- | --- |
| 1 | Poor | High | Vegetation cover <100%<br>or stem density <0.2cm<sup>-2</sup> |
| 2 | Moderate | Moderate | Vegetation cover 100%<br>Stem density >0.2cm<sup>-2</sup><br>Aerial root mat depth <0.5cm |
| 3 | Good | Low | Vegetation cover 100%<br>Stem density >0.2cm<sup>-2</sup><br>Aerial root mat depth >0.5cm |

**Figure S1.**
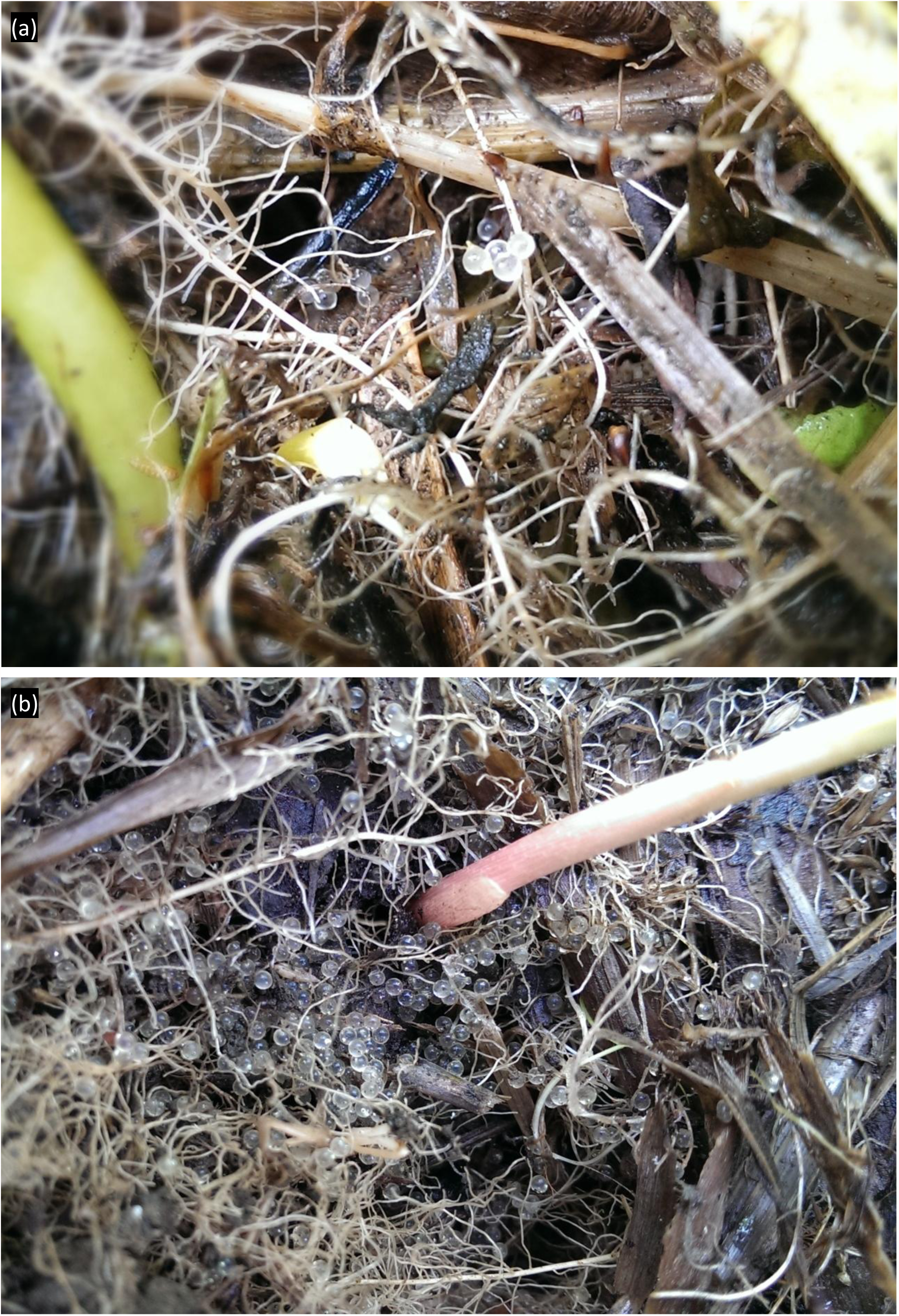
(a) *G. maculatus* eggs laid in riparian vegetation in the Heathcote River, February 2015. Each egg is approximately 1 mm in diameter. (b) An example of high egg densities at the Avondale site in March 2016.

**Figure S2.**
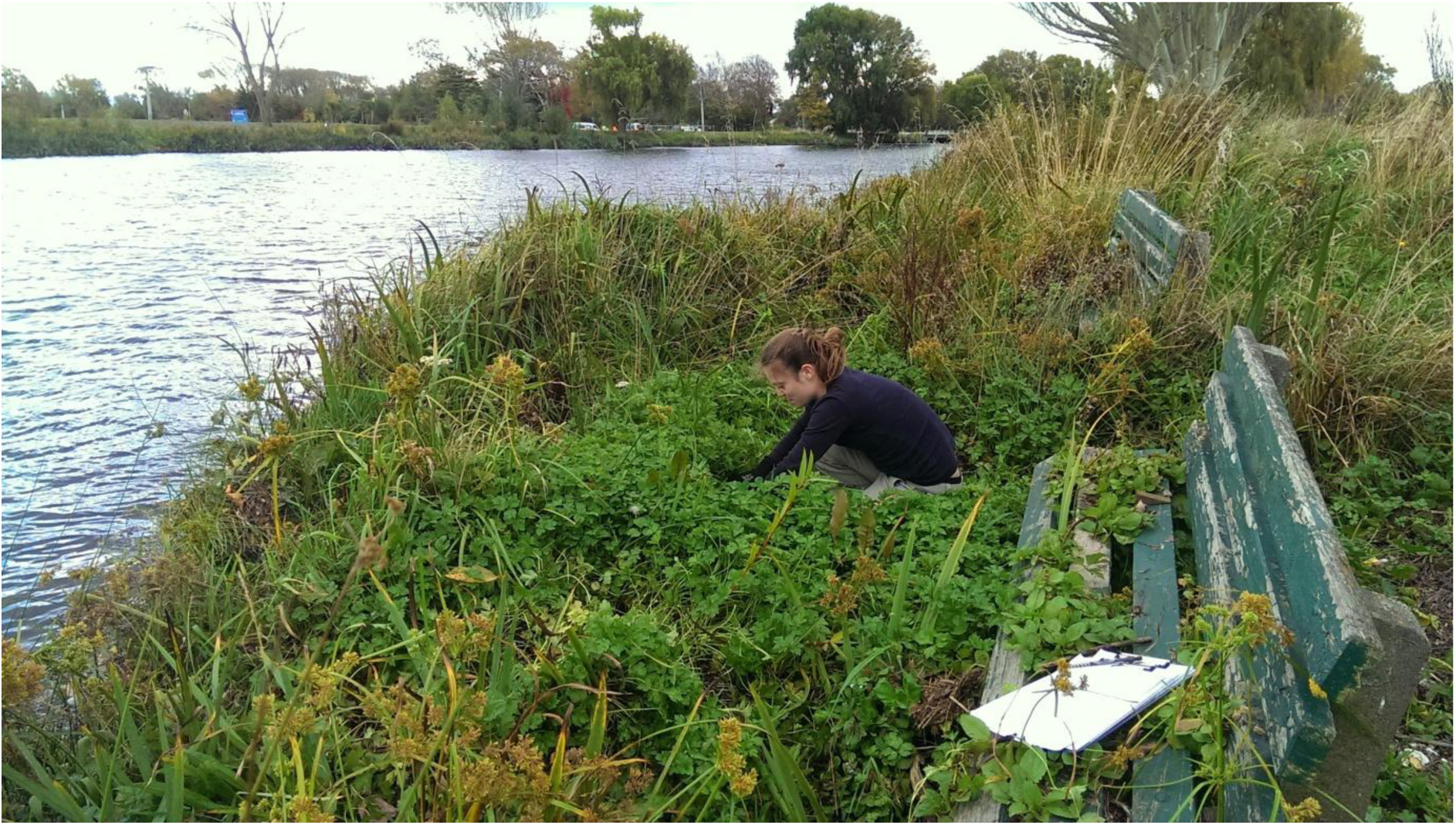
An example of good quality spawning habitat in the Avon River. In this part of the catchment ground levels had dropped by approximately 0.5 m as result of earthquake-induced subsidence and lateral spread. Prior to the earthquakes these overgrown park benches were considerably closers to the waters’ edge.

**Figure S3.**
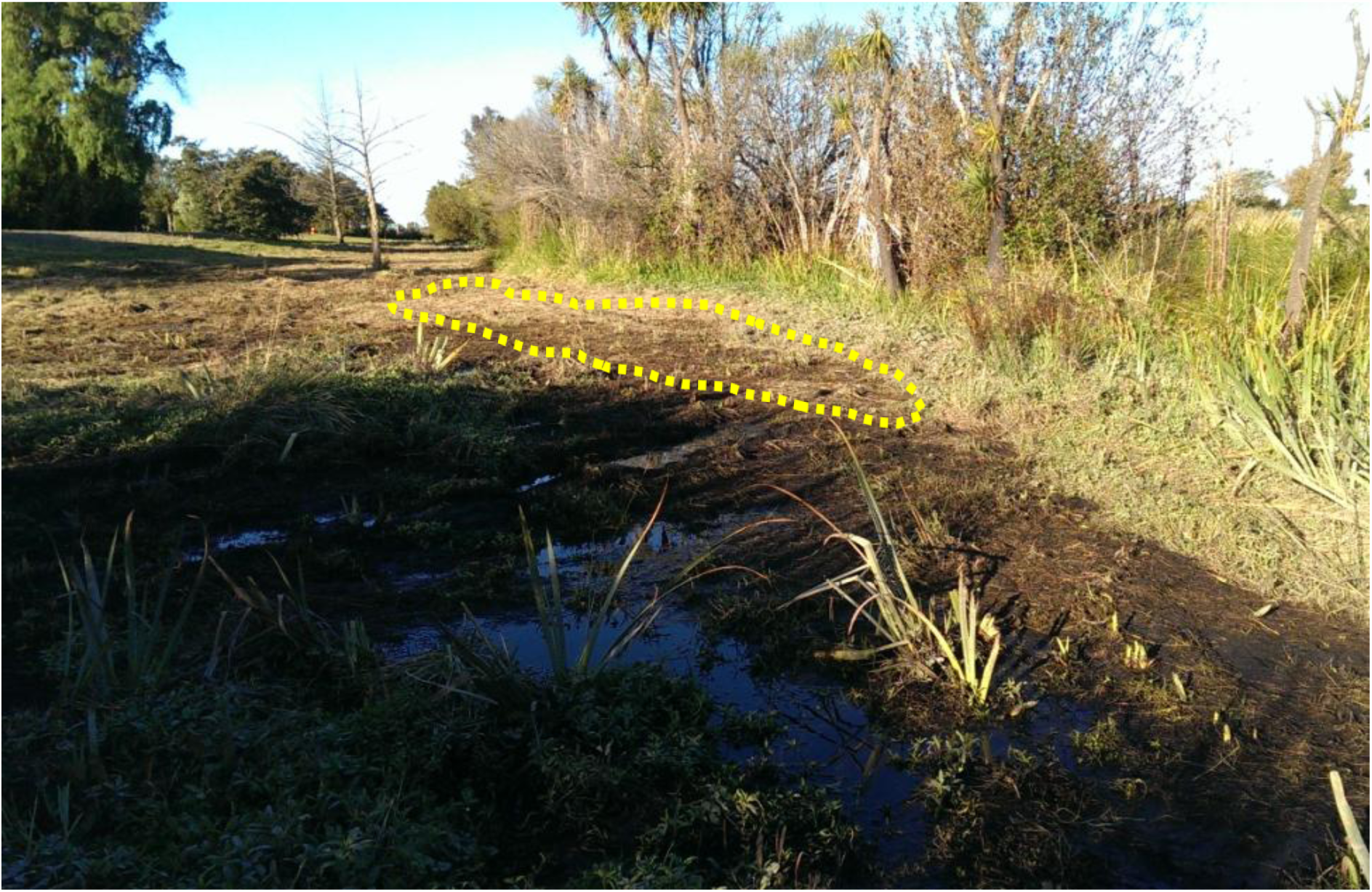
Example of a spawning site affected by mowing in a recreation reserve in the Avon catchment. The dotted line shows the area of occupancy (AOO) prior to mowing in March 2015. Egg mortality is close to 100% in these situations.

**Figure S4.**
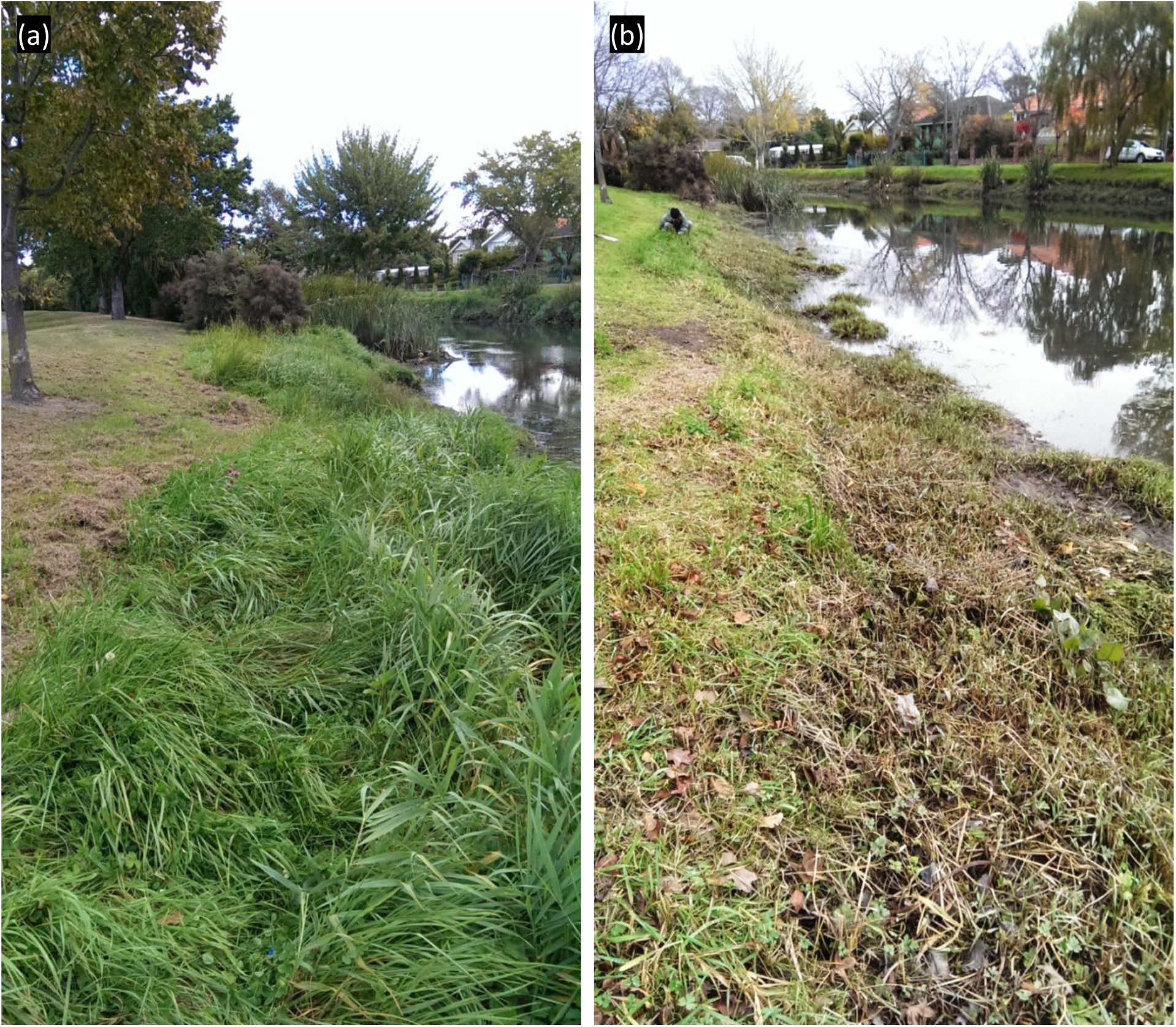
Example of vegetation clearance for flood management. (a) Spawning site in the Heathcote River in March 2015 at which 118, 000 eggs were present (located in the long grass). Mowing of a recreation reserve can also be seen in this image but did not affect the majority of the spawning site which was located lower on the bank face. (b) The same site in early May 2015 showing typical conditions following clearance of vegetation for flood management using line trimmers. This management regime is regularly applied to a large proportion of the study area.

